# Systems analysis by mass cytometry identifies susceptibility of latent HIV-infected T cells to targeting of p38 and mTOR pathways

**DOI:** 10.1101/371922

**Authors:** Linda E. Fong, Victor L. Bass, Serena Spudich, Kathryn Miller-Jensen

## Abstract

Efforts to cure HIV are hindered by viral persistence in latently infected memory CD4+ T cells. Targeting T cell death pathways dysregulated by HIV infection offers a novel approach for eradication of the latent reservoir. To identify potential therapeutic targets, we compared signaling and apoptosis in uninfected and latently infected primary cultured CD4+ central memory T cells by mass cytometry following T cell receptor stimulation. We found that HIV-infected cells were sensitized to activation of pro-apoptotic p38 kinase signaling via p53, and to inhibition of anti-apoptotic mTOR kinase signaling, even without HIV protein expression. Simultaneous targeting of p38 and mTOR kinases in resting CD4+ T cells from virally-suppressed HIV+ patients *ex vivo* reduced cell-associated HIV RNA and DNA. Our results demonstrate how systems biology approaches are useful for identifying novel therapeutic approaches to treat HIV latency, and further suggest that it may be possible to deplete latent HIV-infected T cells without viral reactivation.

## Introduction

HIV proviruses persist in long-lived CD4+ T cells despite suppressive anti-retroviral therapy (ART), and this reservoir of latent virus is a significant obstacle to viral eradication in patients (1-3). Clinical trials with latency reversing agents (LRAs) have successfully activated viral transcription *in vivo* but have shown limited or no depletion of the reservoir (4-7). While many strategies promote the elimination of virus-expressing cells via the immune system (8-11), another approach is to directly target and kill latent HIV-infected T cells (12). There is evidence that HIV infection affects both pro-apoptotic and anti-apoptotic signaling pathways (13-15) and targeting these pathways in combination with LRAs has the potential to reduce reservoir size (16). However, due to the interconnected, multivariate nature of signaling pathways that regulate apoptosis (17-19) and the heterogeneous nature of latent HIV activation (20, 21), successful identification of novel therapeutic targets requires systems-level single-cell analysis.

Mass cytometry provides unprecedented resolution in the multi-dimensional characterization of cellular biomarkers and signaling networks in single cells (22-24). By relying on mass spectrometric time-of-flight (TOF) sampling rather than fluorescence-based detection, mass cytometry enables highly-multiplexed protein detection. This network-level approach has been used to map T cell signaling and functional diversity across several immunological contexts, including pathogen response, hematopoiesis, acute myeloid leukemia, and type 1 diabetes (24-28). Combined with computational techniques to visualize and quantify biological relationships, we can rationally design strategies to target signaling pathways dysregulated in latently infected T cells and enable pharmacological manipulation in a selective manner.

We used mass cytometry to investigate how signaling pathways that regulate apoptosis are altered by latent and reactivating HIV in primary human central memory CD4+ T cells (T_CM_). We computationally reconstructed signal-response functions across uninfected and HIV-infected subpopulations and identified molecular targets that increase apoptosis in latently infected cells without viral reactivation. Finally, we identified clinically-relevant therapeutic agents to target these pathways and demonstrated efficacy in resting CD4+ T cells from HIV-infected patients on suppressive ART. Overall, our study motivates targeted killing of latent HIV-infected T cells as an effective viral eradication strategy.

## Results

### Single-cell mass cytometry reveals increased cell death, despite similar activation dynamics, in primary HIV-exposed T_CM_ cells

We previously showed that TCR stimulation via CD3/CD28 leads to more cell death in latent HIV-infected CD4+ central memory-like T cells infected *in vitro* relative to uninfected cells (29). To investigate systems-level changes in intracellular signaling that explain this observation, we quantified TCR-induced signaling dynamics and apoptosis in uninfected and HIV-infected cells by mass cytometry (Fig. 1A). Primary T cells were differentiated under non-polarizing conditions to induce a central memory T cell (T_CM_) state. On day 5, cells were infected *de novo* with a replication-defective *env(*-) virus construct (DHIV-NEF^+^), pseudotyped with the CXCR4-tropic *env* from HIV-1LAI (30, 31), which generates a mixed population of virus-exposed T_CM_ cells that are uninfected, latently infected, or virus-expressing. Uninfected T_CM_ cells were cultured in parallel for direct comparison (Fig. S1A). After one week in culture, both uninfected and virus-exposed T cells expressed similar levels of memory-like surface markers, and the frequency of virus-exposed cells with basal virus expression was < 10% (Fig. S1B-C).

**Fig. 1.**
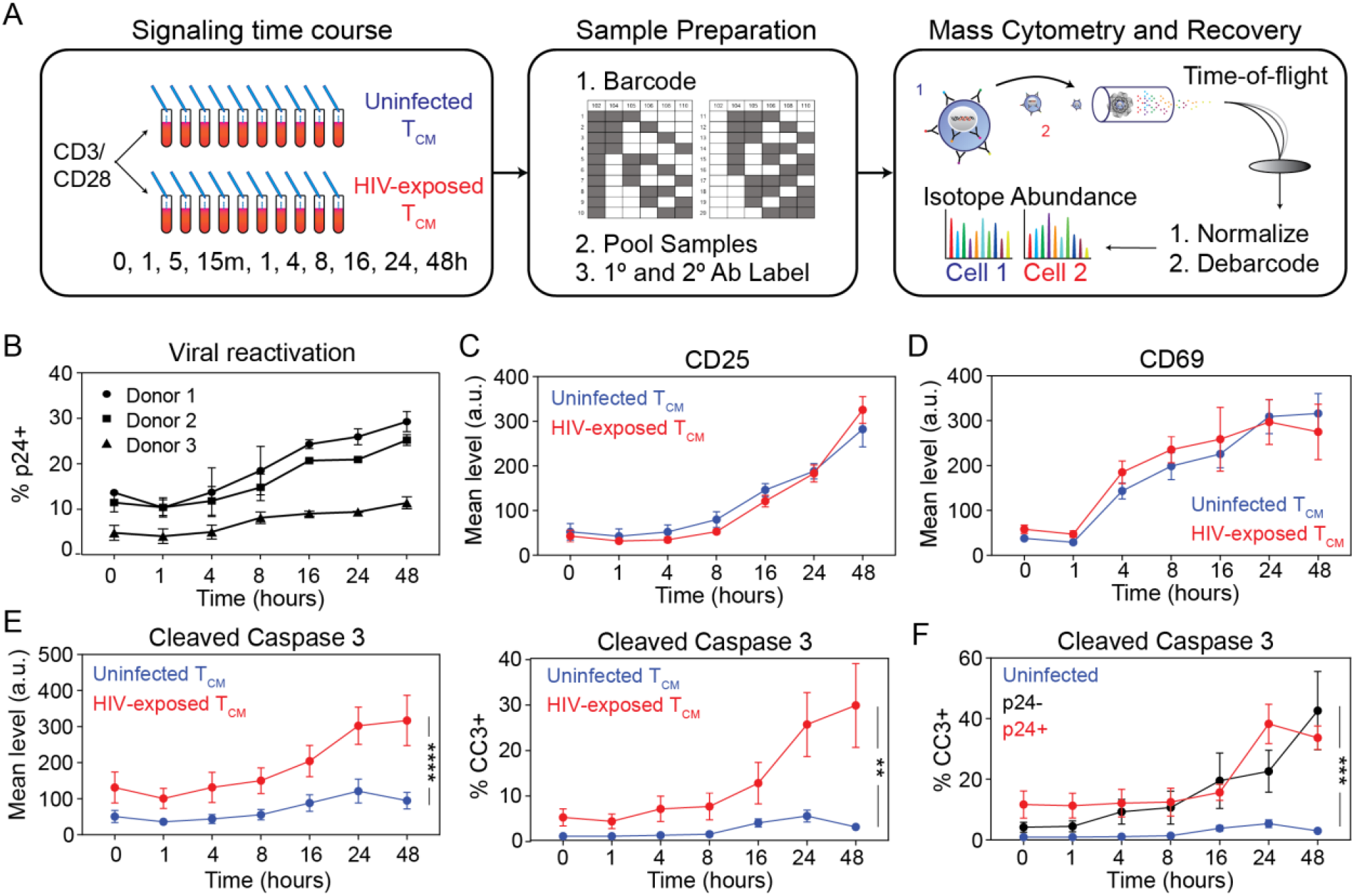
HIV-exposed T_CM_ cells show similar T cell activation but increased apoptosis relative to uninfected cells. (A) Schematic of experimental design. Uninfected and HIV-exposed CD4+ T_CM_ cells were treated in parallel with CD3/CD28. Samples were collected at multiple time points (left), barcoded and pooled (middle), and analyzed by mass cytometry (right). (B) Time course of HIV-exposed T_CM_ cells with reactivating virus (% p24+) by donor following CD3/CD28 activation. Data are presented as means ± SEM for biological replicates. (C-D) Time course of T cell activation markers CD25 (C) and CD69 (D) following CD3/CD28 activation over 48 hours for primary uninfected (blue) and virus-exposed (red) cultured T_CM_ cells. (E) Time course of apoptosis as measured by cleaved caspase 3 (CC3) level and % for the conditions in (C-D). (F) Time course of T_CM_ cells expressing CC3 (%) over 48 hours for uninfected (blue), p24-(black), and p24+ (red) T_CM_. Data are presented as means ± SEM for biological replicates for 3 donors (C-F). Significance calculated over time using a two-way ANOVA (****, p ≤ 0.0001; ***, p ≤ 0.001; **, p ≤ 0.01; *, p ≤ 0.05).

We quantified TCR-induced signaling and apoptosis in uninfected and virus-exposed T_CM_ cells over 48 hours in the presence of ART. Our mass cytometry panel consisted of 6 surface markers, 23 intracellular proteins, and two viral proteins (Table S1). Sampling captured both the shorter timescale of TCR-proximal signaling events (1, 5, 15 and 60 minutes) and the longer timescale of viral reactivation and cell death (4, 8, 16, 24, and 48 hours). To identify virus-expressing cells, we labeled two viral proteins, p24-Gag and Tat. Viral expression in virus-exposed T_CM_ cells after CD3/CD28 stimulation varied from 10%-30% across three donors, as measured by p24-Gag (Fig. 1C), consistent with conventional flow cytometry measurements (Fig. S1C-E). Tat protein levels also increased with time, but the Tat signal was much weaker (Fig. S1F-G), and therefore we used p24-Gag to distinguish between virus-exposed T_CM_ cells with active viral expression and cells that were exposed to virus but remained latent or uninfected.

Virus-exposed T_CM_ cells and uninfected cells showed no difference in T cell activation as measured by the acquisition of surface expression of CD69 and CD25 over 48 hours (Fig. 1C-D). Despite similar T cell-activation dynamics, virus-exposed T_CM_ cells exhibited increased cleaved caspase 3 (CC3), with approximately 30% of virus-exposed cells undergoing apoptosis (i.e., %CC3+) as compared to < 10% of uninfected cells at 48 hours (Fig. 1E). Importantly, the increase in apoptosis was not dependent on the expression of HIV proteins, as the percentage of cells expressing CC3 was similar for p24-and p24+ cells throughout the time course (Fig. 1F). Thus, the dysregulation of apoptosis in virus-exposed T_CM_ cells following CD3/CD28 stimulation appeared to be linked to viral infection but not viral reactivation.

### Virus-exposed T_CM_ cells exhibit systems-level changes in TCR-induced signaling even without HIV protein expression

To identify the signaling networks responsible for differences in apoptosis, we measured the levels of phosphorylated or total protein across multiple pathways activated upon TCR engagement. These pathways included proximal signaling nodes (e.g., p-ZAP70, p-SLP76), kinase signaling pathways (e.g., p-ERK1/2, p-p38), transcription factors (e.g., p-NF-kB, p-STAT1), and apoptosis-related proteins (e.g., p-Bad, p53) (Fig. 2A).

**Fig. 2.**
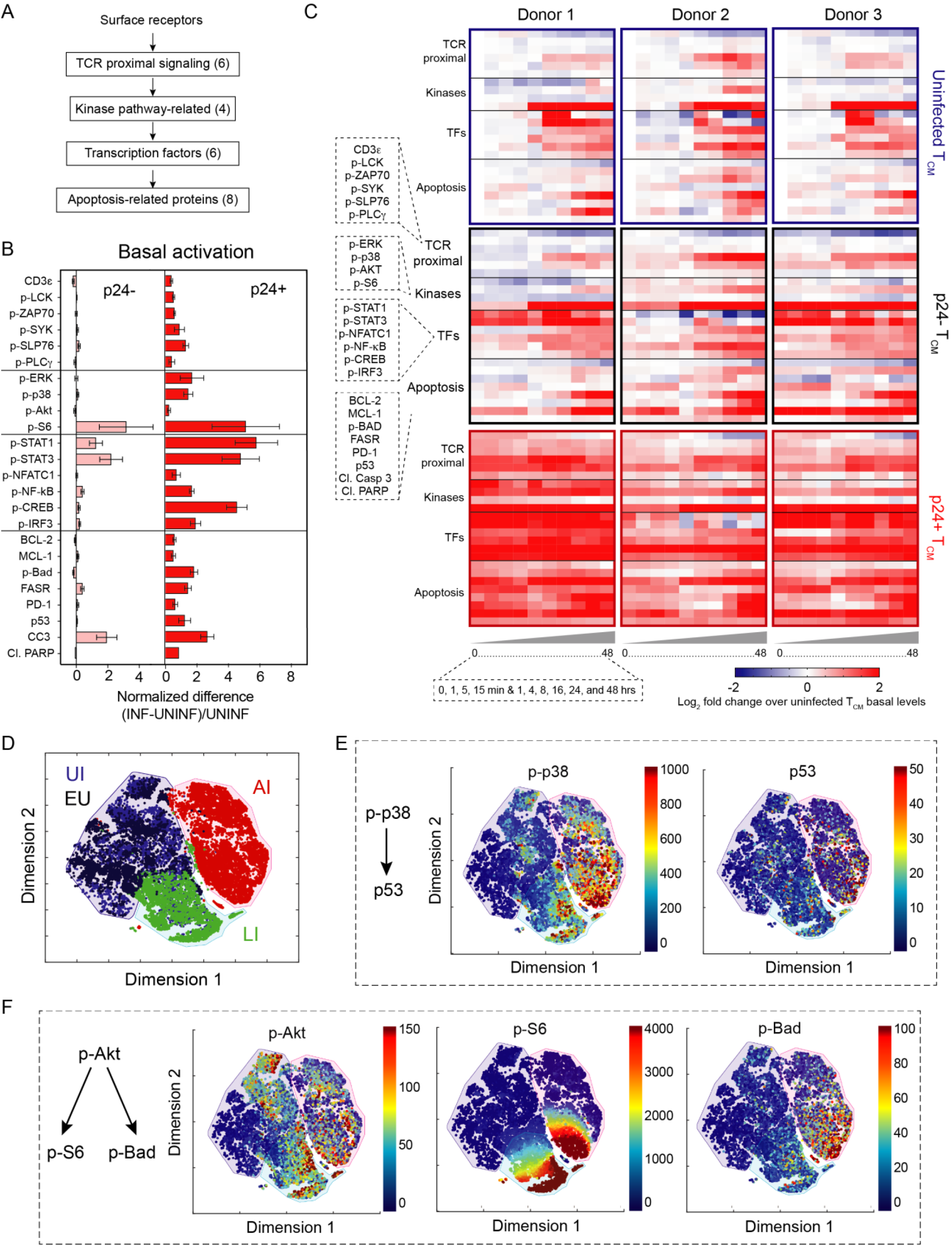
Virus-exposed p24-and p24+ cells exhibit elevated phosphorylation activity and increased expression of apoptotic proteins following TCR stimulation. (A) Overview of signaling and apoptosis proteins measured by mass cytometry (24 total) measured in human primary CD4+ T_CM_ cells from 3 donors. (B) Comparison of basal protein levels in p24-(pink) and p24+ (red) subpopulations relative to uninfected cells. Differences in infected mean abundance values were calculated and normalized to uninfected mean abundance values. Bar graph depicts means ± SEM for 3 donors. *, p ≤ 0.05 by Student’s t-test. (C) Mass cytometry mean abundance values following CD3/CD28 activation for signaling proteins (including phospho-proteins and transcription factors) were measured in uninfected (blue), p24-(black), and p24+ (red) T_CM_ cells over 48 hours for 3 donors. Heatmaps represent the log-2-fold change of the mean mass cytometry abundance values. (D) t-SNE visualization of three cell clusters. Cells in cluster 1 include uninfected (UI) and virus-exposed uninfected (EU) cells; cells in cluster 2 are enriched for latently infected (LI) cells; and cells in cluster 3 are actively infected (AI; see also Fig. S2B-D). (E and F) t-SNE plots for stress signals p-p38 and p53 (E), and for survival signals p-Akt, p-S6 and p-Bad (F) show relative protein expression levels in single cells across clusters.

We separated virus-exposed T_CM_ cells into p24-Gag-and p24-Gag+ subpopulations (referred to as p24-and p24+ cells, respectively), and compared their signaling states to that of uninfected cells prior to CD3/CD28 stimulation. Prior to stimulation, virus-expressing p24+ cells (~5-10% of the virus-exposed population) showed significantly increased phosphorylation levels for MAP kinases (p-ERK and p-p38), p-S6, p-Bad, and several transcription factors (p-STAT1, p-STAT3, p-NF-kB, and p-CREB) (Fig. 2B). By contrast, basal signaling in p24-cells was similar to uninfected cells. Notable exceptions included p-S6, p-STAT1, and p-STAT3, which were significantly increased in p24-cells relative to uninfected T_CM_ cells prior to stimulation (Fig. 2B) and remained elevated following TCR stimulation (Fig. S2A). While increased p-STAT1 and p-STAT3 have been previously observed in cells actively infected with HIV (32, 33), the reason for elevated STAT1/3 activation in p24-cells is unclear.

Following TCR stimulation, we observed systems-level differences in signaling in both p24- and p24+ virus-exposed T_CM_ cells relative to uninfected cells, and these differences were largely conserved across donors (Fig. 2C). p24-cells showed slightly elevated phosphorylation levels for select TCR proximal proteins (p-ZAP70, p-SLP76) within 15 minutes, and this increase appeared to propagate through the MAPK signaling cascades (p-ERK and p-p38). As a result, downstream proteins and transcription factors were more rapidly and strongly induced in p24-cells than in uninfected T_CM_ cells. Signaling in virus-expressing p24+ cells was markedly increased relative to p24-and uninfected T_CM_ cells across almost all measured targets, including TCR proximal and distal signals, demonstrating that the expression of viral proteins significantly affects signaling across multiple pathways in response to TCR stimulation.

We used t-distributed stochastic neighbor embedding (t-SNE) to visualize the relationships between uninfected and virus-exposed cells across all time points, while preserving their local, high-dimensional geometry as determined by the multi-parameter dataset (26, 34). The expression of p24 separated actively infected T_CM_ cells, while uninfected T_CM_ cells overlapped with the virus-exposed p24-cells, consistent with the fact that these cells comprise a mixed population of latently-infected and uninfected cells (Fig. S2B). To separate the overlapping uninfected and virus-exposed p24-cells, we automatically assigned clusters using a nearest-neighbors clustering algorithm (Fig. S2C) (27). By taking the intersection of these two methods, we labeled four phenotypic subpopulations: uninfected (UI), exposed uninfected (EU), latently infected (LI), and actively infected (AI) (Fig. 2D and Fig. S2D). Notably, the UI and EU populations overlapped, while LI and AI populations formed separate clusters.

We visualized the expression of all measured proteins in single cells across clusters (Fig. S2E). Levels of phospho-p38 and its downstream target p53 were increased in the LI and AI populations relative to UI and EU populations (Fig. 2E). We also observed that p-S6 was significantly increased in the LI and AI populations, and p-Bad, the inactive form of the pro-apoptotic mitochondrial Bcl-2 associated death promoter protein Bad, was increased more in the AI population, although there was no corresponding in p-Akt, a shared upstream kinase for S6 and Bad (Fig. 2F). We hypothesized that these observed differences in p38-p53 activation and S6-Bad activation might be related to changes in apoptosis induction. Therefore, we proceeded to examine these pathways to identify and validate new targets for inducing cell death in HIV-infected cells.

### Pro-apoptotic activity of p-p38 and p53 is enhanced in virus-exposed p24-and p24+ T_CM_ cells

We first examined regulation of apoptosis by the p38 MAPK pathway. Although we observed increased average levels of p-p38 and its pro-apoptotic substrate p53 in p24+ cells at all time points, there was no significant increase in p-p38 or p53 levels in p24-cells (Fig. S3 A and B), suggesting that increased abundance alone cannot account for elevated TCR-induced apoptosis. Recent studies suggest that disease states affect interactions between proteins more than protein levels (35). To analyze interactions between proteins and identify how these interactions change between uninfected, p24-and p24+ cell populations, we used conditional density rescaling of protein levels across single cells to visualize the functional relationship between two proteins (36). To quantify signal transmission strength between two nodes along a pathway (e.g. X and Y), we fit a curve to the visualized relationship and calculate the area-under-the-curve (AUC) as previously described (36). The AUC estimates the signal transfer activity between X and Y, or the extent to which an increase in protein X is associated with an increase in protein Y. Differences in AUC across populations (e.g., uninfected and p24+ cells) would suggest an underlying change in the regulation of Y, either directly through X or indirectly via other signaling molecules that also affect Y.

We inferred signal transfer activity (AUC) along the TCR–SLP76–p38–p53–cleaved caspase 3 pathway for uninfected, p24-, and p24+ subpopulations (Fig. 3A and B). Upon CD3/CD28 stimulation, there was no evidence for increased signal transfer activity from p-SLP76 to p-p38 at any time point (Fig. S3C and D). However, between p-p38 and p53, we observed that similar increases in p-p38 were associated with significantly greater increases in p53 in both p24-and p24+ cells relative to uninfected T_CM_ cells at 24 and 48 hours (Fig. 3C).

**Fig. 3.**
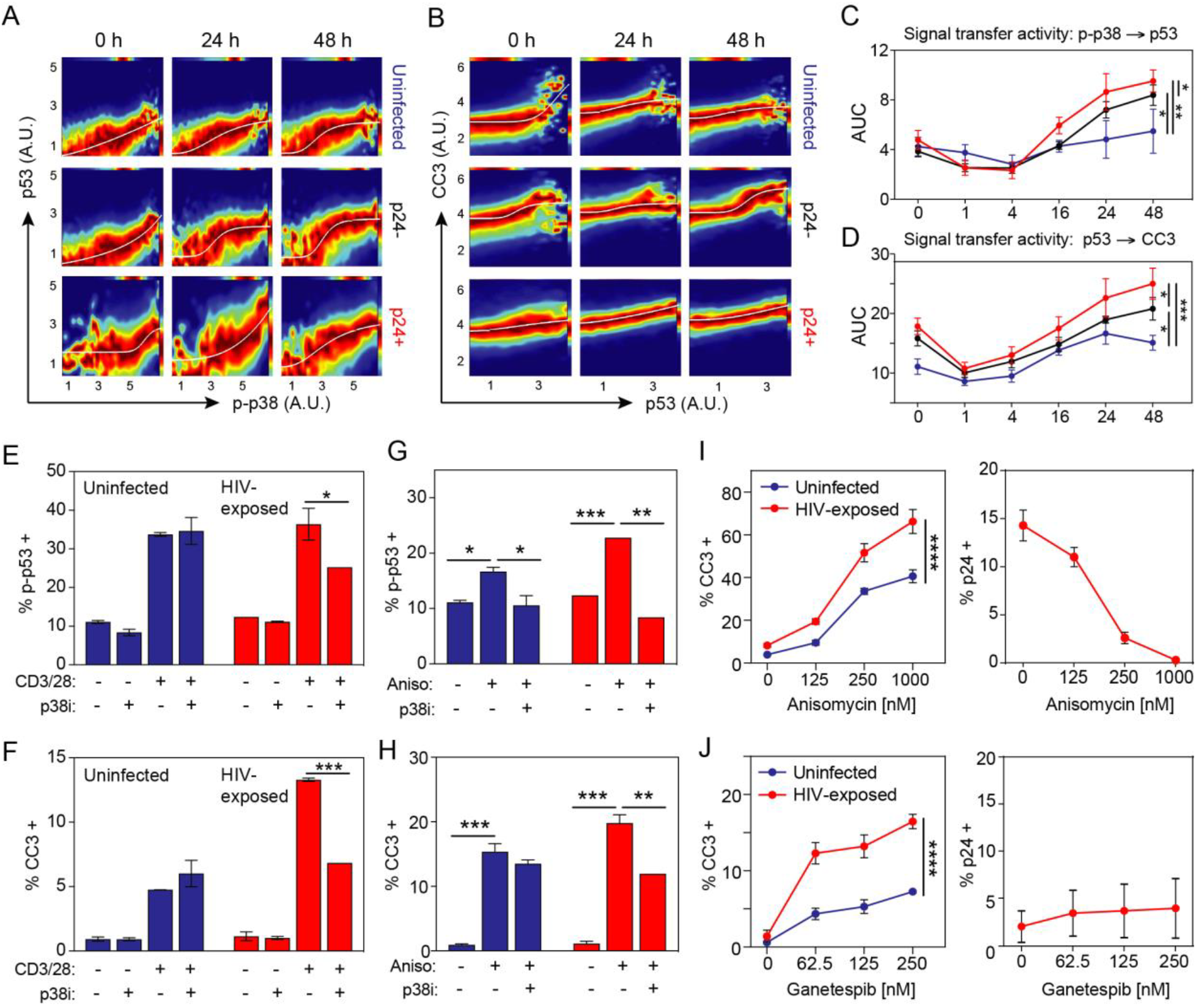
HIV-exposed p24-and p24+ cells exhibit altered apoptotic regulation along the p38-p53-cleaved caspase 3 pathway. (A and B) DREVI plots depict the relationship between p-p38 to p53 (A) and p53 to cleaved caspase 3 (CC3) (B) at 0, 24, and 48 hours. (C and D) Area-under-the-curve (AUC) values for the p-p38-p53 interaction (C) and the p53-CC3 interaction (D) from 0 to 48 hours for uninfected (blue), p24-(black) and p24+ (red) T_CM_ cells. AUC data presented as mean ± SEM for 3 donors. (E) Percentage of p-p53+ cells measured by flow cytometry at 1h with and without p38 inhibition by SB 203580 in the basal state and following CD3/CD28 stimulation in uninfected (blue) and HIV-exposed (red) T_CM_ cells. (F) Percentage of CC3+ cells measured by flow cytometry at 24h with and without SB 203580 in the basal state and following CD3/CD28 stimulation in uninfected (blue) and HIV-exposed (red) T_CM_ cells. (G and H) Percentage of p-p53+ (G) and CC3+ (H) cells measured by flow cytometry with and without SB 203580 following anisomycin stimulation [125nM]. (I and J) Dose-dependent apoptosis (% CC3+; left) and viral reactivation (% p24+; right) following treatment with anisomycin (I) or ganetespib (J) in uninfected (blue) and HIV-exposed (red) T_CM_ cells. Data presented as mean ± SEM for 2 donors. Significance calculated over time or dose using a two-way ANOVA (****, p ≤ 0.0001; ***, p ≤ 0.001; **, p ≤ 0.01; *, p ≤ 0.05).

Signal transfer activity between p53 and CC3 appeared elevated in p24-and p24+ cells prior to stimulation (Fig. 3D), suggesting that accumulation of CC3 occurs at lower thresholds of p53 in virus-exposed T_CM_ subsets. The signal transfer activity between p53 and CC3 increased further following stimulation by CD3/CD28, as demonstrated by higher AUC scores in p24-and p24+ subsets relative to uninfected T_CM_ cells (Fig. 3D). Taken together, we find that the same level of p-p38 is ultimately associated with more CC3 in p24-and p24+ T_CM_ cells by 24-48 hours. Thus, signaling along the p38-p53-CC3 pathway, downstream of p-SLP76, appears to be significantly dysregulated in virus-exposed T_CM_ populations, independent of HIV protein expression.

To validate differential regulation of p53 and cleaved caspase 3 by p-p38, we stimulated uninfected and virus-exposed T_CM_ cells with CD3/CD28 in the presence of the p38 inhibitor SB 203580 and measured the change in p-p53 at 1 hour and CC3 at 24 hours, relative to an uninhibited control. Phosphorylation of p53 at serine 15 attenuates its interaction with negative regulator MDM2, which stabilizes p53 and causes it to accumulate, ultimately leading to growth arrest and apoptosis (37). We found that CD3/CD28 stimulation increased p-p53 to similar levels in uninfected and virus-exposed cells, but p38 inhibition specifically reduced p-p53 only in virus-exposed cells (Fig. 3E). We further found that p38 inhibition significantly reduced CD3/CD28-induced apoptosis (i.e., %CC3+) in virus-exposed cells, but not in uninfected cells (Fig. 3F). These results confirm a regulatory change in p38-mediated apoptosis in latent and reactivating cells that is associated with increased p53.

We recently reported that the antibiotic anisomycin, a potent activator of the p38 and JNK MAPK pathways, increased both p-p38 and p-JNK activation within 30 minutes and cell death at 24 hours in virus-exposed T_CM_ cells relative to uninfected cells (29). We found that anisomycin treatment increased levels of p-p53 and CC3 to a greater extent in virus-exposed cells relative to uninfected cells, and that this could be reversed with p38 inhibition (Fig. 3G-H). Altogether, we conclude that there is increased signal transfer activity along the p38-p53-CC3 axis in latent and virus-expressing cells, which specifically increased their sensitivity to stimulation with CD3/CD28 and to p38 pathway targeting.

Although anisomycin induces more apoptosis in virus-exposed cells than uninfected cells in the absence of viral reactivation (Fig. 3I), the overall toxicity of anisomycin is high, limiting its use as a targeted therapeutic. To reduce non-specific toxicity, we explored small molecules in clinical development that act on p38. We identified ganetespib, a Phase III HSP90 inhibitor that activates p38 when the chaperone HSP90 is sufficiently suppressed, causing apoptotic cell death (38). Similar to anisomycin, ganetespib induced a significantly higher percentage of apoptotic cells in virus-exposed versus uninfected T_CM_ populations in a dose-dependent manner independent of viral reactivation (Fig. 3J). Thus, therapeutic targeting the p38 MAPK pathway has the potential to specifically boost apoptotic signaling in latently infected CD4+ T cells without reactivating latent virus.

### Virus-exposed p24-and p24+ T_CM_ cells exhibit altered Akt-S6 and Akt-Bad signaling and sensitivity to mTOR inhibition

We next examined regulation of the pro-apoptotic protein Bad and the ribosomal protein S6, which share upstream effectors including Akt and 70-kDa ribosomal protein S6 kinase (p70S6K). Akt regulates phosphorylation of S6 via mTORC1-mediated activation of p70S6K, while both Akt and p70S6K are capable of directly phosphorylating Bad and suppressing its apoptotic activity (39-41).

We evaluated the pairwise relationship between p-Akt and p-S6 by visualizing the conditional density-rescaled single-cell data (Fig. 4A). Prior to stimulation, low p-Akt levels were associated with low p-S6 in both uninfected and virus-exposed T_CM_ cells, but moderate levels of p-Akt yielded higher p-S6 in virus-exposed cells (Fig. 4B). Within 4 hours of CD3/CD28 stimulation, signal transfer activity between p-Akt and p-S6 increased similarly in uninfected and virus-exposed T_CM_ populations; but by 8 hours, p24-and p24+ cells exhibited a sustained increase while uninfected T_CM_ cells showed a decrease in activity, as quantified by AUC scores.

**Fig. 4.**
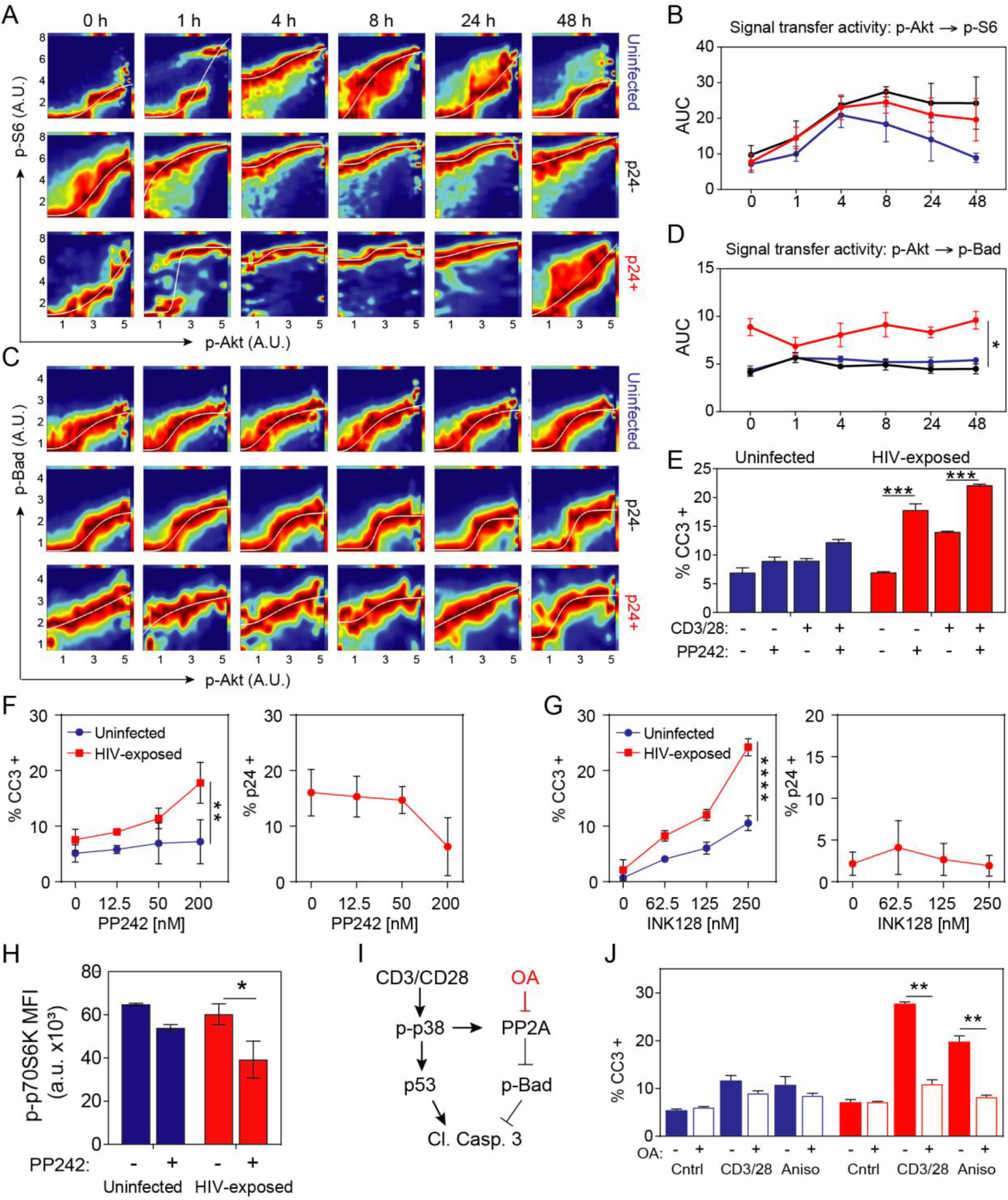
HIV-exposed p24-and p24+ cells show differential activation along the Akt-S6 and Akt-Bad signaling pathways and sensitivity to mTOR inhibition. (A and C) DREVI plots depict the relationship between p-Akt to p-S6 (A) and p-Akt to p-Bad (C) over time (t = 0, 1, 4, 8, 24, 48 h). (B and D) Area-under-the-curve (AUC) values between p-Akt to p-S6 (B) and p-Akt to p-Bad (D) from 0 to 48 hours for uninfected (blue), p24-(black), and p24+ (red) cells. AUC data presented as mean ± SEM for 3 donors. (E) Percentage of cleaved caspase 3+ (CC3+) cells measured by flow cytometry at 24h with and without mTOR kinase inhibition by PP242 [200nM] in uninfected (blue) and HIV-exposed (red) T_CM_ cells. (F-G) Dose-dependent apoptosis (% CC3+; left) and viral reactivation (% p24+; right) following treatment with PP242 (F) and INK128 (G) in uninfected (blue) and HIV-exposed (red) T_CM_ cells. Data presented as mean ± SEM for 2 donors. Significance calculated using a two-way ANOVA (****, p ≤ 0.0001; ***, p ≤ 0.001; **, p ≤ 0.01; *, p ≤ 0.05). (H) Percentage of p-p70S6K+ cells measured by flow cytometry at 1h with and without PP242 [200nM] in uninfected (blue) and HIV-exposed (red) T_CM_ cells. (I) Schematic of p38-Bad crosstalk via PP2A. (J) Percentage of CC3+ cells at 24h with and without okadaic acid [200nM] following stimulation in uninfected (blue) and HIV-exposed (red) T_CM_ cells. **, p ≤ 0.01; *, ≤ 0.05 by Student’s t-test.

When we examined signal transfer activity between p-Akt and p-Bad, we found that virus-expressing p24+ cells exhibited a stronger edge-response function at all time points (Fig. 4C and D), with p-Bad present even at the lowest levels of p-Akt. In contrast, p24-cells showed lower signal transfer activity relative to p24+ cells, suggesting that viral proteins may influence signaling between p-Akt and p-Bad across basal and activated conditions.

Akt activates pro-survival signals (42). Thus, because both p24-and p24+ cells exhibited elevated Akt activity following TCR stimulation, as determined by increased signal transfer activity to p-S6 and p-Bad, we hypothesized that inhibiting a shared upstream kinase would block these pro-survival signals and enhance apoptosis in HIV-infected cells. To block signaling along the Akt-S6 and Akt-Bad pathways, we used the mTOR kinase inhibitor PP242, which competitively inhibits mTOR complexes 1 and 2, thereby reducing the activity of Akt and p70S6K, the kinases directly upstream of Bad and S6. PP242 treatment alone and in combination with CD3/CD28 increased apoptosis (%CC3+) in virus-exposed T_CM_ cells but not uninfected cells at 24 hours (Fig. 4E). Furthermore, PP242 treatment alone induced apoptosis in a dose-dependent manner, even while reducing HIV expression as shown in previous studies (43) (Fig. 4F). A Phase II mTOR kinase inhibitor and PP242 analog INK128, which is in trials for advanced solid tumors and leukemias (44-46), also increased apoptosis in virus-exposed T_CM_ cells without reactivation (Fig. 4G). PP242-induced apoptosis was associated with a significant decrease in p-p70S6K, the kinase downstream of mTOR that phosphorylates S6 and Bad that was specific to HIV-exposed cells (Fig. 4H). Together, these results demonstrate that targeting dysregulated Akt signaling via mTOR inhibition can increase apoptosis in HIV-infected cultured primary CD4+ T_CM_ cells.

We considered why the increased signal transfer activity between Akt and S6 was not also seen between Akt and Bad in latent (p24-) cells. It was previously reported that p38 mediates the dephosphorylation of p-Bad via phosphatase PP2A (47, 48), thereby activating the pro-apoptotic activity of Bad (Fig. 4I). To test whether the lack of increased signal transfer activity between p-Akt and p-Bad in p24-cells might be linked to increased p-p38 activity, we inhibited PP2A with okadaic acid (OA) and measured the change in apoptosis (%CC3+) upon CD3/CD28 stimulation in uninfected and virus-exposed T_CM_ cells. Inhibition of PP2A with OA rescued virus-exposed T_CM_ cells from apoptosis induced by CD3/CD28 but did not affect uninfected cells (Fig. 4J). OA also selectively reversed anisomycin-induced apoptosis in virus-exposed T_CM_ cells. We confirmed this selective rescue by OA in J-DHIV cells, a monoclonal Jurkat-derived cell line with a single latent proviral integration (Fig. S4). Our findings suggest that enhanced PP2A phosphatase activity, perhaps in response to increased p-p38 signaling, may reduce p-Bad levels in latent cells and contribute to the observed loss in signal transfer activity.

### Targeting p38 and mTOR kinase pathways *ex vivo* depletes cell-associated HIV DNA and inducible RNA in resting CD4+ T cells from virally suppressed patients

p38 activation and mTOR inhibition separately increased apoptosis in virus-exposed cells relative to uninfected cells, and therefore we reasoned that targeting these pathways simultaneously might boost cell death in latent CD4+ T cells while minimizing off-target effects in uninfected cells (Fig. 5A). Using the T_CM_ model, we compared cell death in uninfected, p24-, and p24+ T_CM_ cells in response to treatment with ganetespib, INK128, and the combination (Gan+INK) over 48 hours. All treatments significantly increased apoptosis in the virus-exposed p24-population (Fig. 5B), similar to anisomycin, PP242, and anisomycin+PP242 (Fig. S5A). Although ganetespib alone induced the most apoptosis in p24-cells at 48 hours, the combination of Gan+INK accelerated the death rate to induce greater apoptosis at 24 hours. We further tested J-DHIV cells and observed that combining treatments specifically increased apoptosis in latent (GFP-) cells in a dose-dependent manner (Fig. 5C and Fig. S5B). Together these results suggest that combining agents may improve latent HIV-infected cell targeting and increase death kinetics for maximal efficacy (49).

**Fig. 5.**
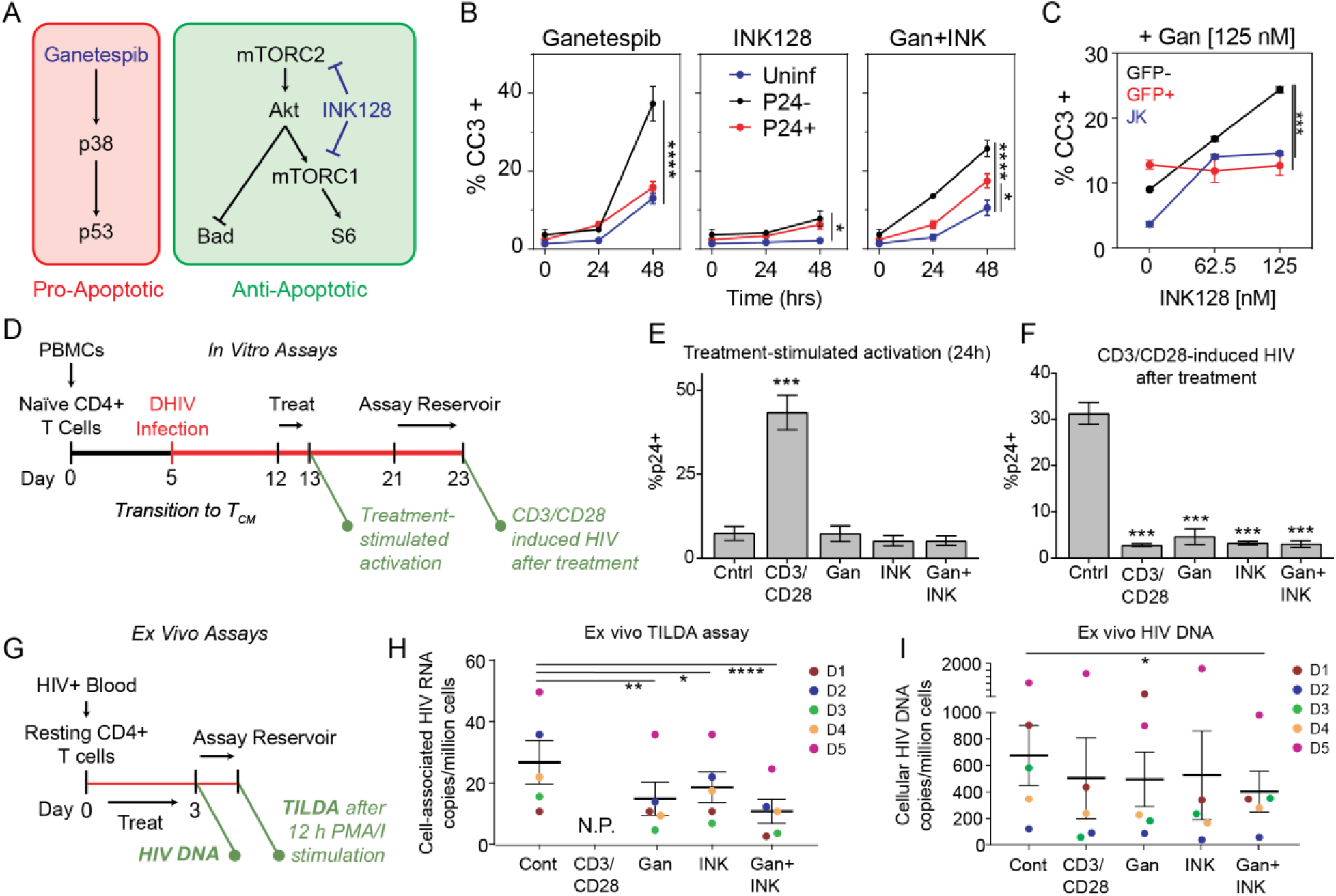
Combinatorial targeting of p38 and mTOR kinase pathways enhances cell death and depletes HIV-infected cells *ex vivo*. (A) Schematic of targeted pathways. (B) Percentage of CC3+ cells measured by flow cytometry following indicated treatment (125nM ganetespib, 125nM INK 128) at 24 and 48 hours. (C) Uninfected Jurkat (blue), GFP– (black) or GFP+ (red) subpopulations of J-DHIV cells that express CC3 at 24h hours by flow cytometry after treatment with 125 nM ganetespib and increasing doses of PP242. Significance calculated over time or dose using a two-way ANOVA (****, p ≤ 0.0001; ***, p ≤ 0.001; **, p ≤ 0.01; *, p ≤ 0.05). (D) Schematic of assay design for measuring HIV-infected cell depletion *ex vivo* in primary cultured T_CM_ cells. (E-F) Viral reactivation (% p24+) induced by treatment with 1:1 CD3/CD28 beads, 125nM ganetespib, 125nM INK128 or a combination measured at 48h by flow cytometry (E) and viral reactivation (% p24+) induced by CD3/CD28 stimulation one week following treatments (F). Data presented as means ± SEM for 2 donors. (G) Schematic of assay design for measuring HIV-infected cell depletion *ex vivo* in resting CD4+ T cells from virally-suppressed patients. (H-I) Cell-associated HIV RNA measured by TILDA assay (53) (H) and cellular HIV DNA copies per million *ex vivo* assayed by nested qPCR (I). Data presented as means ± SEM for 5 HIV+ ART-suppressed donors. Significance calculated using one-way ANOVA, Dunnett’s multiple comparison test (****, p ≤ 0.0001; ***, p ≤ 0.001; **, p ≤ 0.01; *, p ≤ 0.05). N.P., not possible to conduct TILDA due to pre-existing HIV mRNA induced upon initial treatment.

The availability of the small molecules INK128 and ganetespib for inhibiting mTOR and activating p38, respectively, allowed us to explore the potential use of these inhibitors for therapeutic approaches to reduce persistence of virally infected cells. Virus-exposed T_CM_ cells were treated for 24 hours and then cultured post-treatment for an additional week to clear dying cells (Fig. 5D). Then we stimulated all populations with CD3/CD28 to elicit maximal viral reactivation and quantified the extent to which the treatment reduced the percentage of HIV-infected cells relative to a no-treatment control. Treatment with CD3/CD28, which produces near-maximal HIV activation in vitro (50), was included as a positive control. We found that CD3/CD28 induced significant viral expression by 24 hours, while ganetespib and INK128, alone or in combination, did not (Fig. 5E). Despite this lack of viral activation, ganetespib, INK128 and the combination all significantly reduced the percentage of latent-but-inducible HIV-infected cells to nearly the same extent as CD3/CD28 (Fig. 5F). Similar results were observed for anisomycin and PP242 (Fig. S5C-D). To determine if this reduction also corresponded with a reduction in cell-associated HIV DNA, we measured proviral DNA copies before and after treatment (Fig. S5E). We found that treatment with p38-or mTORC-targeted therapies also moderately decreased average HIV cell-associated DNA levels, with combinatorial treatments of anisomycin+PP242 or Gan+INK128 exhibiting approximately a 30% reduction (Fig. S5E).

Finally, we tested whether targeting the p38 and mTOR pathways would reduce inducible RNA in resting CD4^+^ T cells and cell-associated HIV DNA from virally suppressed HIV+ subjects on ART. Resting CD4+ T cells were isolated from patient whole blood and treated for 72 hours *ex vivo* with ganetespib, INK128, or the combination (Fig. 5H). We reasoned that if ganetespib and/or INK128 preferentially induced apoptosis in latent HIV-infected cells, then cell-associated RNA and DNA should be reduced following treatment. To measure inducible cell-associated HIV RNA remaining after treatment, we used a Tat/Rev inducible limiting dilution assay (TILDA) (51), in which a population of cells from each treatment group was stimulated with PMA+ionomycin (PMA/I) for 12 hours and inducible cell-associated HIV RNA was quantified by qPCR. Genomic DNA was collected from the remaining cells and cell-associated HIV DNA remaining after treatment was quantified by nested qPCR. We used CD3/CD28 as a benchmark for cell-associated HIV DNA depletion (52-56), but the cell-associated HIV transcription activated by CD3/CD28 during the initial 3-day treatment prevented its use as a benchmark in the TILDA assay.

Ganetespib, INK128, and Gan+INK all significantly reduced the level of PMA/I-induced cell-associated RNA relative to the untreated control, with Gan+INK causing the largest decrease of >50% (Fig. 5I). Furthermore, HIV DNA levels relative to an untreated control were reduced with CD3/CD28, ganetespib, INK128, and combinatorial treatment with Gan+INK, with Gan+INK resulting in a significant reduction of 33% (Fig 5J). These findings suggest that combinatorial treatment with Gan+INK targets and clears resting CD4+ cells harboring inducible proviruses in the absence of viral reactivation. Overall, these Phase II/III drugs show promise as therapeutic agents for targeted depletion of latent HIV-infected cells in ART-suppressed patients.

## Discussion

Through systems analyses of high-dimensional single-cell data, we demonstrated that latent and reactivating HIV alters apoptosis regulation in primary cultured T_CM_ cells. Virus-exposed p24- and p24+ T_CM_ cells exhibited increased signaling activity along the p38-p53-CC3 pathway and were sensitized to p38 pathway targeting with anisomycin and ganetespib (Fig. 3). Our findings are consistent with transcriptomic RNA-Seq measurements demonstrating upregulation of p53-related genes upon latency reversal (57). We further showed that inhibition of mTOR signaling also increased apoptosis in latently infected cells (Fig. 4). Importantly, previous work found that first-generation mTOR inhibitors, like rapamycin, downregulated pro-inflammatory markers while maintaining CTL recognition ability for infected cell killing (58). Inhibition of mTOR signaling has also been demonstrated to promote maintenance of latency (43). Here we add the new observation that mTOR may be activating anti-apoptotic signals in latent-HIV infected cells that can be targeted for clearance. Finally, we showed that targeting these pathways in combination *ex vivo* reduced cell-associated HIV DNA and RNA in resting CD4+ T cells isolated from ART-suppressed HIV+ patients (Fig. 5).

This study identifies targetable differences in signaling regulation to eliminate latent HIV-infected cells without viral reactivation. Although there is evidence that HIV protein expression is cytotoxic (59, 60) and marks host cells as targets of cytotoxic T cells, HIV proteins can also contribute to pro-survival signaling (61, 62). Recently, cellular survival programs governed by BIRC5, a molecular inhibitor of apoptosis, were shown to support long-term survival of HIV-infected CD4+ T cells (63). Moreover, the Bcl-2 family has long been implicated in balancing low-level HIV virus production with cell survival (64-67). We further showed that reactivating infected (p24+) cells express higher levels of anti-apoptotic p-Bad than latently infected (p24-) cells, suggesting that activation of viral protein expression may complicate eradication efforts.

Looking towards translational studies, we identified a Phase III HSP90 inhibitor targeting p38 and a Phase II mTOR kinase inhibitor that effectively enhanced apoptosis in latently infected CD4+ T_CM_ cells. Our systems approach revealed novel categories of therapeutic agents that, unlike existing LRAs, act to directly eliminate latent HIV-infected T cells. HSP90 inhibitors have been in clinical trials since 2005 (68), with next-generation agents continuously being developed (69). Several PP242 analogs have also entered clinical trials, including INK128, OSI-027, AZD8055, and AZD2014 (70). Although the small molecules tested in this study likely exhibit some pleiotropic effects, advances in drug development and delivery, in addition to further treatment optimization, will likely reduce these effects. For example, different HSP90 and mTOR inhibitors have been shown to exhibit varying rates of apoptosis (49), and therefore it may be possible to systematically assess timing of therapeutic administration to improve efforts for *in vivo* eradication. Overall, our findings support a new clinical strategy aimed at directly killing latently-infected CD4+ T cells that complements other strategies for latency eradication.

## Materials and Methods

### Participants with HIV on ART

All participants positive for HIV were on combination ART and had had undetectable plasma viral loads (<50 copies/ml) for at least 1 year (median 11 years) (Supplementary Table 2). HIV+ subjects were recruited as described in Protocol No. 1502015318: HIV Associated Neurocognitive Disorder and Neurocognitive Disorders with Other Infectious Diseases, and the study was approved by the Yale Human Investigation Committee. All research participants gave written informed consent.

### Resting CD4+ T cells from HIV+ patients

100 mL of whole blood was collected from HIV+ donors under a Yale-approved HIC protocol to investigate anti-latency targets. Peripheral blood mononuclear cells (PBMCs) were harvested by density centrifugation with Ficoll-Paque (GE Healthcare). CD4+ T cells were isolated from the PBMC population using the STEMCELL EasySep™ Human CD4 + T Cell Enrichment Kit, (19052, negative selection), according to manufacturer specifications. Resting CD4+ T cells were then negatively selected for by removing CD69+, CD25+ and HLA-DR+ subsets using Miltenyi Microbead kits (CD69 Microbead Kit II, human, 130-092-355; CD25-Biotin Microbeads, human, 130-091-235, Anti-HLA-DR Microbeads, human, 130-046-101) and LS columns, according to manufacturer specifications.

### Virus production

The DHIV-Nef+ plasmid (containing a small out-of-frame deletion in the gp120-coding area that renders it defective in Env), DHIV-GFP plasmid (DHIV plasmid containing GFP in place of Nef) and the pLET-LAI plasmid (encoding an X4 envelope gene) were kind gifts from Dr. Vincente Planelles. The envelope gene was originally derived from a CXCR4-dependent isolate, HIV-1-LAI, and the *env* glycoprotein was produced with the expression construct pLET-LAI (31, 71, 72). To generate virus, DHIV-Nef+ and DHIV-GFP were co-transfected with pLET-LAI plasmids into HEK293T cells (gift from Dr. David Schaffer) using standard calcium phosphate-mediated transfection. After 18 h, the transfection medium was replaced with 10 ml of fresh medium (IMDM + 10% FBS and L-glutamine in the absence of antibiotics, all from Gibco Life Technologies) and cultured for an additional 36 h. Supernatants were then collected and pre-cleared by centrifugation. After centrifugation, the supernatant was filtered, aliquoted, and frozen at −80 °C. To normalize infections, p24 was quantified in virus-containing supernatants by enzyme-linked immunosorbent assay (ELISA; ZeptoMetrix).

### Primary cultured central memory CD4+ T cells (T_CM_) activation and infection

Buffy coats of leukocytes were obtained from anonymous HIV-donors (New York Blood Center). Peripheral blood mononuclear cells (PBMCs) were harvested by density centrifugation with Ficoll-Paque (GE Healthcare). Naïve CD4^+^ T cells were isolated by MACS negative selection using the human naïve T-cell isolation kit (Miltenyi Biotec) to yield a population with the phenotype CD4^+^ CD45RA^+^ CD45RO^−^ CCR7^+^ CD62L^+^ CD27^+^ with purity levels equal or higher than 95%. Cells were then activated for 72 hours with beads coated with αCD3 and αCD28 antibodies, Dynabeads Human T-Activator CD3/CD28 for T cell Expansion and Activation, (ThermoFisher Scientific) under non-polarizing (NP) conditions, in the presence of 10 ng/ml of TGF-β1, 2 μg/ml of anti-Human IL-12 and 1 μg/ml of anti-Human IL-4 (all from Peprotech). Dynabeads were removed on day 3 and consequently cultured at a density of 1 × 10^6^ cells/ml in complete medium with 30 IU/ml of rhIL-2 (R&D Systems). Cells were infected by spinoculation on Day 5: 10^6^ cells were infected with 300 ng/mL p24 for 2 hours at 2900 rpm and 37°C in 1 mL. After infection, cells kept in culture at 1 × 10^6^ cells/ml in complete medium with 30 IU/ml of rhIL-2 and 10 μM indinavir sulfate (ART, protease inhibitor) at 37 °C for 7 days prior to further experimentation, during which actively infected cells die, leaving a mixed population of latently infected or uninfected cells.

### Jurkat T cell culture and infection (J-DHIV cell line)

Jurkat cells clone E6-1 (ATCC) were cultured in RPMI media (Gibco Life Technologies) supplemented with 10% FBS (Atlanta Biologicals) and penicillin and streptomycin (Gibco Life Technologies at a concentration of 2×10^5^-2×10^6^ cells/ml at 37°C and 5% CO_2_. Cells were infected with a DHIV-GFP/X4 virus and GFP-cells were sorted to isolate uninfected and latently infected populations. Cells were then stimulated with TNFα (Peprotech), and GFP+ cells were sorted into a 96-well plate to establish a clonal line of latently infected cells(29). This parental Jurkat and J-DHIV clonal cell line model provides a fully infected population with minimal divergence between uninfected and infected counterparts.

### Mass cytometry time course collection, sample preparation and processing

All stimulations were conducted in the presence of 10 μM indinavir sulfate. Cells treated with CD3/CD28 were collected 5 minutes prior to the expiration of the time point and placed on a magnet for bead removal. Cells were treated for 1 minute with Cisplatin (25mM) to distinguish dead cells. Cells were fixed by adding ice-cold PFA to a final concentration of 2% PFA and then fixed for 30 minutes at 4°C. To ensure the detection of subtle differences in cellular parameters over time (i.e. shifts in protein phosphorylation upon stimulation), we utilized a pooled sample analysis approach to minimize sample-to-sample variation. A binary labeling strategy composed of 6 palladium-based barcodes was used to label samples prior to immune-staining with a single antibody cocktail (73). Cells were washed 1x with MAXPAR staining buffer after fixation and once with 0.02% saponin in PBS, then barcoded for 30 minutes at room temperature using the Cell-ID 20-Plex Barcoding Kit (Fluidigm) in the presence of 0.02% saponin (Affymetrix 00-8333-56 Permeabilization Buffer, 10X). After measurement, barcode combinations were identified with a doublet-filtering, deconvolution algorithm and used recover the individual samples for further analysis.

Following barcoding, the samples were washed 2x with MAXPAR cell staining buffer and combined into a single sample. Metal conjugated antibodies were sorted into two groups: surface marker or intracellular and stained consecutively. Samples were incubated with the surface marker cocktail for 45 minutes at room temperature. Samples were washed 2x, re-fixed with 2% PFA, and permeabilized with methanol to aid in the detection of phospho-targets. Samples were then incubated with the intracellular staining cocktail for 45 minutes at room temperature. Samples were washed 2x in MAXPAR staining buffer and once with 2% PFA then incubated for overnight with an iridium-containing DNA intercalator (DVS/Fluidigm) in 2% PFA at 4°C. On the following day, samples were washed twice with MAXPAR staining buffer, and once in water. The cell concentration adjusted to 700,000 cells/ml in water supplemented with 10% EQ 4-element calibration beads (Fluidigm), filtered, and injected successively to a CyTOF II mass cytometer in a series of 500 μl aliquots. At the end on the acquisition, the files resulting from the different injections were concatenated into one .fcs file, and the calibration beads were removed using the CyTOF II built-in software. Detector sensitivity was performed at the beginning and at the end of each use of CyTOF II using polystyrene normalization beads containing lanthanum-139, praseodymium-141, terbium-159, thulium-169, and lutetium-175.

### Mass cytometry data analysis

Mass cytometry mean abundance values were transformed using a scaled arcsinh (scaling factor 5 used to minimize noise near the limit of detection) and represented in heatmaps as the log-2-fold change over the uninfected basal condition. Software for implementing viSNE, Phenograph, and DREVI, and the associated CYT tools were downloaded from the Pe’er Lab website (https://www.c2b2.columbia.edu/danapeerlab/html/software.html). viSNE (26) was implemented using default parameters of the CYT tool. We performed clustering using PhenoGraph (27) on a combined sample of 180,000 cells (subsampling of 2000 cells per time point for uninfected, p24-, and p24+ subgroups across 3 donors) with 5000 neighbors and a Euclidian distance-based metric. Transformed data were analyzed using DREVI as previously described (36).

### Flow cytometry assays

Cells were fixed with and permeabilized with the BD Cytofix/Cytoperm solution (BD Biosciences). Cells were washed and stained in the presence of BD Perm/Wash Buffer (BD Biosciences). For primary cells, viral activation was measured by intracellular staining for p24, an HIV-1 capsid protein, with the monoclonal antibody AG3.0 at a 1:40 dilution (NIH AIDS Reagent Program, courtesy of Dr. Jonathan Allan) and anti-mouse IgG secondary antibodies conjugated to AlexaFluor 647 (ThermoFisher Scientific). For Jurkat latent infections, activation was quantified by analyzing green fluorescent protein (GFP) level in 10,000 cells. Cell death was measured after 10 minutes of staining with propidium iodide (Sigma Aldrich) at a working concentration of 5 μg/ml. Flow cytometry assays were analyzed on an Accuri C6 Cytometer (BD Biosciences).

### Genomic DNA isolation and quantitative PCR to measure *in vitro* HIV DNA

Genomic DNA was extracted using the DNeasy Blood and Tissue kit (Qiagen) per the manufacturer’s specifications with the following modifications: before eluting DNA columns were dried with an additional 1 minute spin at maximum speed, elution volume was 100uL Buffer AE. Total HIV DNA from *in vitro* infections was quantified by qPCR. 3μL of genomic DNA at 50pg/μL was combined with 2μL of either 5μM LTR or CD3 primer mix and 5μL SsoAdvanced Universal SYBR Green Supermix (Bio-Rad). Cycling conditions for both primers were as follows: denaturation for 3 minutes at 98°C, then 40 cycles of denaturation for 15 seconds at 98°C then annealing and extension for 1 minute at 59°C with a fluorescence read each cycle, followed by melt-curve analysis to confirm specificity of primers. LTR forward and reverse primers were designed using Primer-BLAST(74), CD3 primer sequences were from *Vandergeeten et al(75)*. Primer amplicon standard curves were used to calculate starting quantities of CD3 and LTR DNA.

### *Ex vivo* TILDA Assay, RNA Isolation, and RT-PCR to measure cell-associated RNA from ART-suppressed patients

The inducible RNA was measured using the TILDA assay developed by *Procopio et al(51)*. Following drug treatment, CD4 + T cells were serially diluted and plated in a 96 well plate at 50 × 103 cells/well, and stimulated for 12 h with 10 ng/ml PMA and 1 μM Ionomycin (Sigma) in the presence of 10 μM indinavir sulfate (ART, protease inhibitor) and 10 μM Amprenavir (ART, protease inhibitor). After stimulation, cells were lysed in TRIzol reagent. RNA was extracted using Zymo Research Direct-zol™-96 RNA plates (R2056) and eluted into 25 μL of water. 1 μl of the purified RNA was used for RT-PCR and transfered a 96 well plate containing 5 μl of 2 × reaction buffer from the SuperScript III Platinum One-Step qRT-PCR Kit (Life Technologies). Pre-amplification was carried out by adding 5 μl of a PCR mix containing 0.2 μl Superscript III Platinum Taq (Life Technologies), 0.05 μl RNasin Plus RNase Inhibitor (Promega), 0.125 μl of each primer (tat1.4 and rev both at 20 μM), 2.2 μl Tris–EDTA (TE) buffer and 2.3 μl H_2_O to each well (final reaction volume = 11 μl). All primer sequences were adapted from *Pasternak et al*(76). Pre-amplification was carried out using the following steps: reverse transcription at 50 °C for 15 min, denaturation at 95 °C for 2 min, 24 cycles of amplification (95 °C 15 s, 60 °C 4 min) on a CFX Connect Real-Time PCR instrument (Biorad). At the end of the pre-amplification, 40 μl of TE buffer was added to each well and 1 μl of the diluted PCR products was used as template for the *tat2*/*rev* real-time PCR reaction. This reaction was performed by adding 5 μl of the SsoAdvanced Universal SYBR Green Supermix (Bio-Rad), 0.2 μl of each primer (tat2 and rev, both at 20 μM), and 3.6 μl H_2_O to each well (final reaction volume = 10 μl). The real-time PCR reaction was carried out in a CFX Connect Real-Time PCR instrument (BioRad) using the following program: Preincubation 95 °C for 10 min, 40 cycles of amplification (95 °C 10 s, 60 °C 30 s), followed by melt curve analysis to confirm specificity of primers. Positive wells at each dilution were counted and the infectious units per million were calculated using a maximum likelihood estimate and 95% confidence intervals; infection frequency calculator (http://silicianolab.johnshopkins.edu/) (77).

### Nested PCR to measure *ex vivo* HIV DNA from ART-suppressed patients

Total HIV DNA in patient cells was quantified using nested PCR adapted from *Vandergeeten et al (75)*. For the first round of PCR, 15 μl of genomic DNA at 20 ng/μl was used in 50 μl reaction containing 1X ThermoPol buffer (NEB), 1 mM MgSO_4_ (NEB), 300 μM dNTPs (Thermo), 300 nM each of four outer primers (ULF1, UR1, outerCD3_fwd, and outerCD3_rev), and 2.5 U Taq polymerase (NEB). The first round cycling conditions were: a denaturation step of 8 min at 95°C and 12 cycles of amplification (95°C for 1 min, 55°C for 40 s, 72°C for 1 min), followed by a final elongation step at 72°C for 15 min. First round products were diluted 1:10 in water before the second round of PCR. For the second round, 3 μl of diluted product was combined with 2 μl of 5 μM inner primer mix (either LambdaT + UR2 or innerCD3_fwd + innerCD3_rev) and 5 μl of SsoAdvanced Universal SYBR Green Supermix (BioRad). The second round cycling conditions for both primer pairs were: a denaturation step of 4 min at 95°C and 40 cycles of amplification (95°C for 10 s, 60°C for 10 s), followed by melt curve analysis to confirm specificity of primers. This reaction was carried out in a CFX Connect Real-Time PCR instrument (BioRad).

### Primers used in study

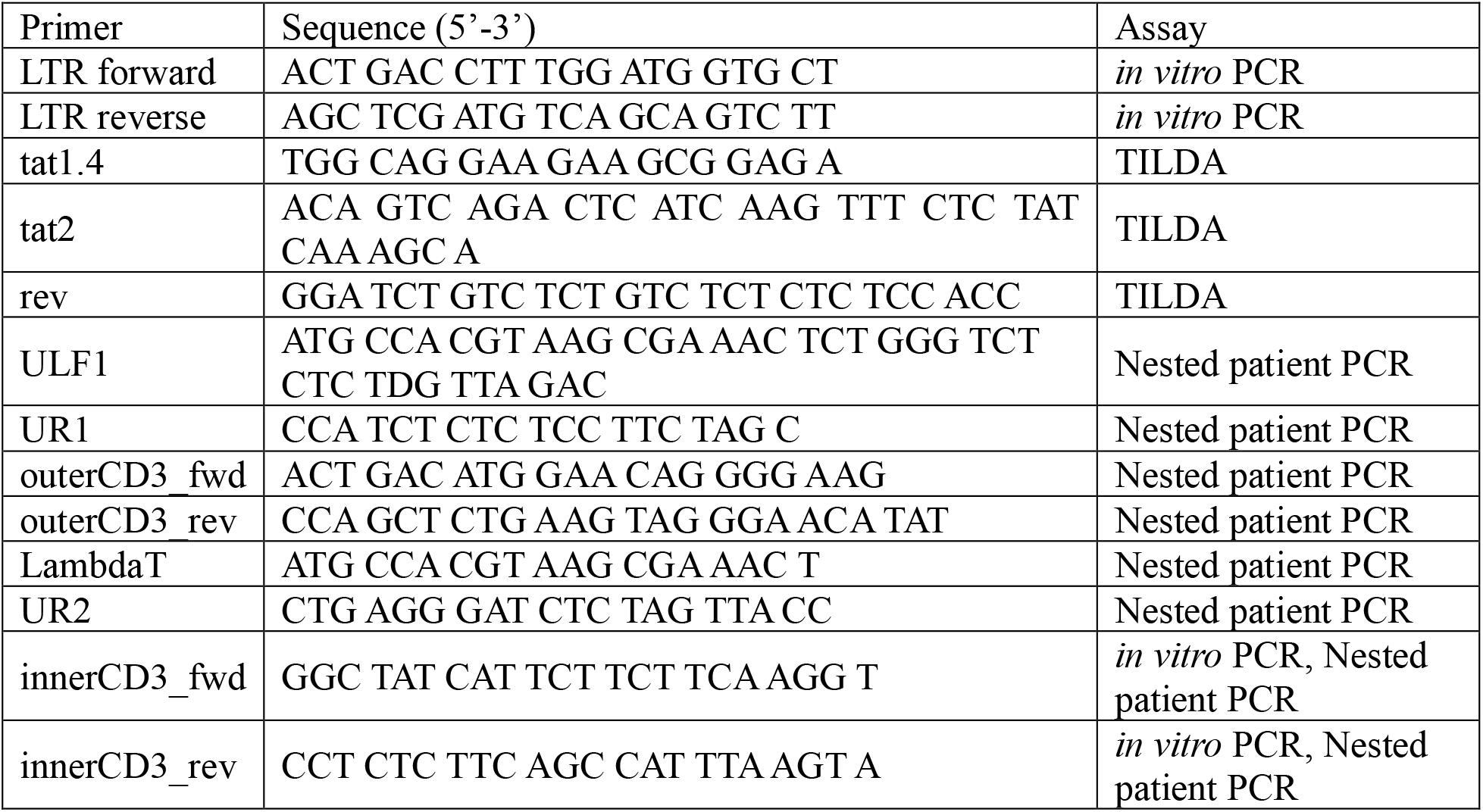

## Acknowledgements

We thank Dr. Bernd Bodenmiller for his advice in optimizing the mass cytometry experiments. We thank Dr. Ruth Montgomery and Shelly Ren at the Yale CyTOF Core for processing the samples and facility support. We thank Dr. Smita Krishnaswamy for providing guidance on DREVI analysis. We thank Dr. Vicente Planelles and Dr. Adam Spivak for their help in designing the *in vitro* HIV DNA assays. We thank Jennifer Chiarella and Tobias Kirchwey for coordinating patient recruitment. We thank Dr. Ya-Chi Ho for her guidance designing the *ex vivo* experiments and for reviewing the manuscript. The following reagent was obtained through the NIH AIDS Reagent Program, Division of AIDS, NIAID, NIH: Indinavir Sulfate and Amprenavir. Funding for this work was provided by NIH 1R21AI132013 (to K.M-J.). V.L.B. was supported by NIH predoctoral training grant in virology (5T32AI055403).

## Author Contributions

L.E.F. and K.M-J. designed the in vitro experiments. L.E.F. collected the *in vitro* data and performed the data analysis. L.E.F. and V.L.B. collected the *ex vivo* HIV+ patient data. S.S. recruited the HIV-infected subjects. K.M-J. and L.E.F. wrote the manuscript. All authors edited the manuscript.

## Additional Information

The authors declare no competing financial interests.

## References

1. Finzi D (1997) Identification of a Reservoir for HIV-1 in Patients on Highly Active Antiretroviral Therapy. Science 278(5341):1295-1300.

2. Chun TW, et al. (1997) Quantification of latent tissue reservoirs and total body viral load in HIV-1 infection. Nature 387.

3. Wong JK, et al. (1997) Recovery of replication-competent HIV despite prolonged suppression of plasma viremia. Science 278.

4. Archin NM, et al. (2012) Administration of vorinostat disrupts HIV-1 latency in patients on antiretroviral therapy. Nature 487(7408):482-485.

5. Archin NM, et al. (2017) Interval dosing with the HDAC inhibitor vorinostat effectively reverses HIV latency. J Clin Invest 127(8):3126-3135.

6. Elliott JH, et al. (2014) Activation of HIV transcription with short-course vorinostat in HIV-infected patients on suppressive antiretroviral therapy. PLoS Pathog 10(10):e1004473.

7. Elliott JH, et al. (2015) Short-term Disulfiram to Reverse Latent HIV Infection: A Phase 2 Dose Escalation Study. The lancet. HIV 2(12):e520-e529.

8. Brockman MA, Jones RB, & Brumme ZL (2015) Challenges and Opportunities for T-Cell-Mediated Strategies to Eliminate HIV Reservoirs. Frontiers in immunology 6:506.

9. Marsden MD & Zack JA (2015) Experimental Approaches for Eliminating Latent HIV. Forum on immunopathological diseases and therapeutics 6(1-2):91-99.

10. Garrido C, et al. (2016) HIV Latency-Reversing Agents Have Diverse Effects on Natural Killer Cell Function. Frontiers in immunology 7:356.

11. Wu G, et al. (2017) HDAC inhibition induces HIV-1 protein and enables immune-based clearance following latency reversal. JCI insight 2(16).

12. Kim Y, Anderson JL, & Lewin SR (Getting the “Kill” into “Shock and Kill”: Strategies to Eliminate Latent HIV. Cell Host & Microbe 23(1):14-26.

13. Timilsina U & Gaur R (2016) Modulation of apoptosis and viral latency - an axis to be well understood for successful cure of human immunodeficiency virus. J Gen Virol 97(4):813-824.

14. Cummins NW & Badley AD (2014) Making sense of how HIV kills infected CD4 T cells: implications for HIV cure. Molecular and cellular therapies 2:20.

15. Fernández Larrosa PN, et al. (2008) Apoptosis resistance in HIV-1 persistently-infected cells is independent of active viral replication and involves modulation of the apoptotic mitochondrial pathway. Retrovirology 5(1):19.

16. Cummins NW, et al. (2016) Prime, Shock, and Kill: Priming CD4 T Cells from HIV Patients with a BCL-2 Antagonist before HIV Reactivation Reduces HIV Reservoir Size. J Virol 90(8):4032-4048.

17. Benedict KF & Lauffenburger DA (2013) Insights into Proteomic Immune Cell Signaling and Communication via Data-Driven Modeling. Systems Biology, ed Katze MG (Springer Berlin Heidelberg, Berlin, Heidelberg), pp 201-233.

18. Badley AD, Sainski A, Wightman F, & Lewin SR (2013) Altering cell death pathways as an approach to cure HIV infection. Cell Death Dis 4:e718.

19. McLean JE, Ruck A, Shirazian A, Pooyaei-Mehr F, & Zakeri ZF (2008) Viral manipulation of cell death. Current pharmaceutical design 14(3):198-220.

20. Ho Y-C, et al. (2013) Replication-Competent Noninduced Proviruses in the Latent Reservoir Increase Barrier to HIV-1 Cure. Cell 155(3):540-551.

21. Wong VC, et al. (2018) NF-kappaB-Chromatin Interactions Drive Diverse Phenotypes by Modulating Transcriptional Noise. Cell Rep 22(3):585-599.

22. Bandura DR, et al. (2009) Mass Cytometry: Technique for Real Time Single Cell Multitarget Immunoassay Based on Inductively Coupled Plasma Time-of-Flight Mass Spectrometry. Analytical Chemistry 81(16):6813-6822.

23. Leelatian N, Diggins KE, & Irish JM (2015) Characterizing Phenotypes and Signaling Networks of Single Human Cells by Mass Cytometry. Single Cell Protein Analysis: Methods and Protocols, eds Singh AK & Chandrasekaran A (Springer New York, New York, NY), pp 99-113.

24. Bendall SC, et al. (2011) Single-Cell Mass Cytometry of Differential Immune and Drug Responses Across a Human Hematopoietic Continuum. Science 332(6030):687-696.

25. Newell EW, Sigal N, Bendall SC, Nolan GP, & Davis MM (2012) Cytometry by time-of-flight shows combinatorial cytokine expression and virus-specific cell niches within a continuum of CD8+ T cell phenotypes. Immunity 36(1):142-152.

26. Amir E-aD, et al. (2013) viSNE enables visualization of high dimensional single-cell data and reveals phenotypic heterogeneity of leukemia. Nat Biotech 31(6):545-552.

27. Levine Jacob H, et al. (2015) Data-Driven Phenotypic Dissection of AML Reveals Progenitor-like Cells that Correlate with Prognosis. Cell 162(1):184-197.

28. Mingueneau M, et al. (2014) Single-cell mass cytometry of TCR signaling: Amplification of small initial differences results in low ERK activation in NOD mice. Proceedings of the National Academy of Sciences 111(46):16466-16471.

29. Fong LE, Sulistijo ES, & Miller-Jensen K (2017) Systems analysis of latent HIV reversal reveals altered stress kinase signaling and increased cell death in infected T cells. Scientific Reports 7(1):16179.

30. Bosque A & Planelles V (2009) Induction of HIV-1 latency and reactivation in primary memory CD4+ T cells. Blood 113(1):58-65.

31. Bosque A & Planelles V (2011) Studies of HIV-1 latency in an ex vivo model that uses primary central memory T cells. Methods 53(1):54-61.

32. Kohler JJ, Tuttle DL, Coberley CR, Sleasman JW, & Goodenow MM (2003) Human immunodeficiency virus type 1 (HIV-1) induces activation of multiple STATs in CD4+ cells of lymphocyte or monocyte/macrophage lineages. J Leukoc Biol 73(3):407-416.

33. Venkatachari NJ, et al. (2015) Temporal transcriptional response to latency reversing agents identifies specific factors regulating HIV-1 viral transcriptional switch. Retrovirology 12(1):1-22.

34. van der Maaten LJP & Hinton GE (2008) Visualizing data using t-SNE. Journal of Machine Learning Research 9((Nov)):2431-2456.

35. Zhong Q, et al. (2009) Edgetic perturbation models of human inherited disorders. Mol Syst Biol 5:321.

36. Krishnaswamy S, et al. (2014) Conditional density-based analysis of T cell signaling in single-cell data. Science 346(6213).

37. Dumaz N & Meek DW (1999) Serine15 phosphorylation stimulates p53 transactivation but does not directly influence interaction with HDM2. Embo j 18(24):7002-7010.

38. Ota A, Zhang J, Ping P, Han J, & Wang Y (2010) Specific Regulation of Non-canonical p38α Activation by Hsp90-Cdc37 Chaperone Complex in Cardiomyocyte. Circulation research 106(8):1404-1412.

39. Altomare DA & Khaled AR (2012) Homeostasis and the Importance for a Balance Between AKT/mTOR Activity and Intracellular Signaling. Current Medicinal Chemistry 19(22):3748-3762.

40. Luo J, Manning BD, & Cantley LC (2003) Targeting the PI3K-Akt pathway in human cancer. Cancer Cell 4(4):257-262.

41. Harada H, Andersen JS, Mann M, Terada N, & Korsmeyer SJ (2001) p70S6 kinase signals cell survival as well as growth, inactivating the pro-apoptotic molecule BAD. Proceedings of the National Academy of Sciences of the United States of America 98(17):9666-9670.

42. Lawlor MA & Alessi DR (2001) PKB/Akt: a key mediator of cell proliferation, survival and insulin responses? J Cell Sci 114(Pt 16):2903-2910.

43. Besnard E, et al. (2016) The mTOR Complex Controls HIV Latency. Cell Host & Microbe 20(6):785-797.

44. Gokmen-Polar Y, et al. (2012) Investigational drug MLN0128, a novel TORC1/2 inhibitor, demonstrates potent oral antitumor activity in human breast cancer xenograft models. Breast cancer research and treatment 136(3):673-682.

45. Janes MR, et al. (2012) Efficacy of the investigational mTOR kinase inhibitor MLN0128/INK128 in models of B-cell acute lymphoblastic leukemia. Leukemia 27:586.

46. Benjamin D, Colombi M, Moroni C, & Hall MN (2011) Rapamycin passes the torch: a new generation of mTOR inhibitors. Nature Reviews Drug Discovery 10:868.

47. Grethe S & Porn-Ares MI (2006) p38 MAPK regulates phosphorylation of Bad via PP2A-dependent suppression of the MEK1/2-ERK1/2 survival pathway in TNF-alpha induced endothelial apoptosis. Cellular signalling 18(4):531-540.

48. Junttila MR, Li SP, & Westermarck J (2008) Phosphatase-mediated crosstalk between MAPK signaling pathways in the regulation of cell survival. FASEB journal : official publication of the Federation of American Societies for Experimental Biology 22(4):954-965.

49. Forcina GC, Conlon M, Wells A, Cao JY, & Dixon SJ (2017) Systematic Quantification of Population Cell Death Kinetics in Mammalian Cells. Cell Systems 4(6):600-610.e606.

50. Bosque A, Famiglietti M, Weyrich AS, Goulston C, & Planelles V (2011) Homeostatic Proliferation Fails to Efficiently Reactivate HIV-1 Latently Infected Central Memory CD4+ T Cells. PLOS Pathogens 7(10):e1002288.

51. Procopio FA, et al. (2015) A Novel Assay to Measure the Magnitude of the Inducible Viral Reservoir in HIV-infected Individuals. EBioMedicine 2(8):874-883.

52. Bullen CK, Laird GM, Durand CM, Siliciano JD, & Siliciano RF (2014) New ex vivo approaches distinguish effective and ineffective single agents for reversing HIV-1 latency in vivo. Nat Med 20(4):425-429.

53. Laird GM, et al. (2015) Ex vivo analysis identifies effective HIV-1 latency-reversing drug combinations. J Clin Invest 125(5):1901-1912.

54. Díaz L, et al. (2015) Bryostatin activates HIV-1 latent expression in human astrocytes through a PKC and NF-ĸB-dependent mechanism. Scientific Reports 5:12442.

55. Gutierrez C, et al. (2016) Bryostatin-1 for latent virus reactivation in HIV-infected patients on antiretroviral therapy. AIDS (London, England) 30(9):1385-1392.

56. Mehla R, et al. (2010) Bryostatin Modulates Latent HIV-1 Infection via PKC and AMPK Signaling but Inhibits Acute Infection in a Receptor Independent Manner. PLoS ONE 5(6):e11160.

57. White CH, et al. (2016) Transcriptomic Analysis Implicates the p53 Signaling Pathway in the Establishment of HIV-1 Latency in Central Memory CD4 T Cells in an In Vitro Model. PLOS Pathogens 12(11):e1006026.

58. Cheng Y, Wong MT, van der Maaten L, & Newell EW (2016) Categorical Analysis of Human T Cell Heterogeneity with One-Dimensional Soli-Expression by Nonlinear Stochastic Embedding. Journal of immunology (Baltimore, Md. : 1950) 196(2):924-932.

59. Costin JM (2007) Cytopathic Mechanisms of HIV-1. Virology Journal 4(1):100.

60. Rasheed S, Gottlieb AA, & Garry RF (1986) Cell killing by ultraviolet-inactivated human immunodeficiency virus. Virology 154(2):395-400.

61. Malim MH & Emerman M (2008) HIV-1 Accessory Proteins—Ensuring Viral Survival in a Hostile Environment. Cell Host & Microbe 3(6):388-398.

62. Geleziunas R, Xu W, Takeda K, Ichijo H, & Greene WC (2001) HIV-1 Nef inhibits ASK1-dependent death signalling providing a potential mechanism for protecting the infected host cell. Nature 410(6830):834-838.

63. Kuo H-H, et al. (2018) Anti-apoptotic Protein BIRC5 Maintains Survival of HIV-1-Infected CD4<sup>+</sup> T Cells. Immunity 48(6):1183-1194.e1185.

64. Zhang M, et al. (2002) Bcl-2 upregulation by HIV-1 Tat during infection of primary human macrophages in culture. Journal of biomedical science 9(2):133-139.

65. Aillet F, et al. (1998) Human Immunodeficiency Virus Induces a Dual Regulation of Bcl-2, Resulting in Persistent Infection of CD4(+) T-or Monocytic Cell Lines. Journal of Virology 72(12):9698-9705.

66. Petrovas C, et al. (2004) HIV-specific CD8+ T cells exhibit markedly reduced levels of Bcl-2 and Bcl-xL. Journal of immunology (Baltimore, Md. : 1950) 172(7):4444-4453.

67. Cummins NW, et al. (2017) Maintenance of the HIV reservoir is antagonized by selective BCL2 inhibition. Journal of Virology.

68. Kim YS, et al. (2009) Update on Hsp90 inhibitors in clinical trial. Current topics in medicinal chemistry 9(15):1479-1492.

69. Butler LM, Ferraldeschi R, Armstrong HK, Centenera MM, & Workman P (2015) Maximizing the Therapeutic Potential of Hsp90 Inhibitors. Molecular cancer research : MCR 13(11):1445-1451.

70. Sun S-Y (2013) mTOR kinase inhibitors as potential cancer therapeutic drugs. Cancer Letters 340(1):1-8.

71. Challita-Eid PM, et al. (1998) Inhibition of HIV Type 1 Infection with a RANTES-IgG3 Fusion Protein. AIDS Research and Human Retroviruses 14(18):1617-1624.

72. Andersen JL, et al. (2006) HIV-1 Vpr-Induced Apoptosis Is Cell Cycle Dependent and Requires Bax but Not ANT. PLOS Pathogens 2(12):e127.

73. Zunder ER, et al. (2015) Palladium-based mass tag cell barcoding with a doublet-filtering scheme and single-cell deconvolution algorithm. Nat. Protocols 10(2):316-333.

74. Ye J, et al. (2012) Primer-BLAST: a tool to design target-specific primers for polymerase chain reaction. BMC Bioinformatics 13:134.

75. Vandergeeten C, et al. (2014) Cross-clade ultrasensitive PCR-based assays to measure HIV persistence in large-cohort studies. J Virol 88(21):12385-12396.

76. Pasternak AO, et al. (2008) Highly sensitive methods based on seminested real-time reverse transcription-PCR for quantitation of human immunodeficiency virus type 1 unspliced and multiply spliced RNA and proviral DNA. J Clin Microbiol 46(7):2206-2211.

77. Rosenbloom DIS, et al. (2015) Designing and interpreting limiting dilution assays: general principles and applications to the latent reservoir for HIV-1. bioRxiv.

